# Probabilistic integration of transcriptome-wide association studies and colocalization analysis prioritizes molecular pathways of complex traits

**DOI:** 10.1101/2022.07.19.500651

**Authors:** Jeffrey Okamoto, Lijia Wang, Xianyong Yin, Francesca Luca, Roger Pique-Regi, Adam Helms, Hae Kyung Im, Jean Morrison, Xiaoquan Wen

## Abstract

Transcriptome-wide association studies (TWAS) and colocalization analysis are complementary integrative genetic association approaches routinely used to identify functional units underlying complex traits in post-genome-wide association study (post-GWAS) analyses. Recent studies suggest that both approaches are individually imperfect, but joint usage can yield robust and powerful inference results. This paper introduces a new statistical framework, INTACT, to perform probabilistic integration of TWAS and colocalization evidence for implicating putative causal genes. This procedure is flexible and can work with a wide range of existing TWAS and colocalization approaches. It has the unique ability to quantify the uncertainty of implicated genes, enabling rigorous control of false-positive discoveries. Taking advantage of this highly-desirable feature, we describe an efficient algorithm, INTACT-GSE, for gene set enrichment analysis based on the integrated TWAS and colocalization analysis results. We examine the proposed computational methods and illustrate their improved performance over the existing approaches through simulation studies. Finally, we apply the proposed methods to the GTEx data and a variety of GWAS summary statistics derived from complex and molecular traits previously analyzed by Hukku *et al*. and Sinnott-Armstrong *et al*. We find empirical evidence that the proposed methods improve and complement existing putative gene implication methods and are advantageous in evaluating and identifying key gene sets and biological pathways.

## 1 Introduction

Recent improvements in sequencing technologies have led to numerous genome-wide association studies (GWAS) that reveal variant-level genetic associations underlying complex diseases. However, a lack of understanding of the molecular mechanisms through which these variants act remains a significant barrier to understanding disease etiology and advancement of treatments. Specifically, for many statistically significant GWAS loci, the underlying target genes cannot be confidently identified [1, 2, 3, 4]. Linking genetic associations to putative causal genes (PCGs) remains a long-standing open problem in statistical genetics. Traditionally, PCG identification relies on relevant biological knowledge of genes in the proximity of GWAS loci. Henceforth, we refer to such methods as proximity-plus-knowledge-based approaches [5]. The extent of the existing knowledge base can limit these methods, and they cannot quantify the uncertainty of the PCG implications. With the increasing availability of genome-scale molecular phenotyping, a new class of emerging computational methods has been developed to establish a relationship between genetic variants, molecular phenotypes, and complex traits by integrating GWAS and molecular quantitative trait loci (QTL) data [6, 7, 8, 9, 10, 11, 12, 13, 14, 15]. These methods have shown promise in not only identifying PCGs but also implicating relevant molecular mechanisms. Thus, we refer to this class of methods as mechanism-aware PCG implication approaches. The most commonly used molecular phenotype data for integrative genetic association analysis are currently expression QTL (eQTL) data. Successful applications have been extended to other types of molecular data, including chromatin structure [16, 17, 18], protein [19, 20], and metabolite data [21, 22].

Two popular types of mechanism-aware methods that use transcriptome data for PCG identification are transcriptome-wide association studies (TWAS) and colocalization analyses. TWAS examine correlation between genetically predicted gene expression and a given complex trait [8, 23, 7, 6, 10]. Under the instrumental variable (IV) analysis assumptions, such correlations imply potential causal relationships from gene expressions to traits [24, 6]. Colocalization analysis attempts to identify overlaps between causal GWAS hits and causal QTLs [25, 11, 13]. However, it does not distinguish between the case of vertical pleiotropy (i.e., gene expressions mediating complex traits) and the case of horizontal pleiotropy. Despite many successful applications, both types of approaches exhibit some noticeable limitations. Colocalization analysis is known to have low sensitivity due to the intrinsic difficulty in pinpointing causal genetic variants in the presence of linkage disequilibrium (LD) [26]. TWAS is known to have low specificity due to LD hitchhiking. It is caused by modest to weak LD between predictive eQTLs and causal GWAS variants and can drive spurious TWAS correlation signals without true causal relationships [24, 14, 9, 27]. Depending on the relationship between the causal eQTLs and GWAS hits, LD hitchhiking can occur between genes or within a single gene. Existing approaches, e.g., FOCUS [9], can resolve between-gene LD hitchhiking. However, resolving within-gene LD hitchhiking remains an open problem [14]. Hukku et al. [27] explore the inferential reproducibility of PCG implications between TWAS and colocalization analysis, concluding that these two procedures can be complementary. Particularly, they argue that both TWAS and colocalization signals should be present for a putative causal gene based on theoretical and empirical evidence. In practice, although an *ad hoc* procedure proposed therein has demonstrated the effectiveness of this analytical strategy [10], rigorous computational methods with the ability to quantify evidence and uncertainty for PCGs are still lacking.

Beyond implicating individual PCGs, there is an even greater need to explore underlying biological pathways and gene sets to dissect disease etiology. The problem is often framed in similar scientific contexts as gene set enrichment (GSE) analysis, where relevant gene sets are expected to be over-represented by genes with specific properties. Although GSE analysis seems to be a natural downstream analysis for PCG implications (i.e., by treating putative causality as the gene property of interest), it has not been a common practice in the field. To the best of our knowledge, GSE analysis methods that explicitly account for the uncertainty of implicated PCGs do not exist in the literature.

This paper proposes a Bayesian procedure for probabilistic evidence integration, INTACT (INtegrative TWAS And ColocalizaTion), to identify PCGs using results from a TWAS scan and colocalization analysis. It is built upon the ideas from [27] but preserves uncertainty from the two forms of analysis by systematically combining results into a gene-level probabilistic quantification of putative causality. We show that INTACT is robust to LD hitchhiking and more powerful than the existing colocalization analysis approaches. Furthermore, we adapt ideas from the previous GSE analysis methods [28, 29] to develop a new computational method, INTACT-GSE, which performs PCG enrichment analysis utilizing the probabilistic output from INTACT. We examine the performance of INTACT and INTACT-GSE using simulations and multiple real data sets. Our real data applications involve complex phenotypes from [30, 10], for which a great deal of molecular biology is known. This prior knowledge allows us to compare the performance of our procedure with a proximity-plus-knowledge-based PCG implication method.

We implement our computational methods in an open-source R package, INTACT. It is freely available at https://github.com/jokamoto97/INTACT/.

The code to reproduce our analysis can be found at htpps://github.com/jokamoto97/intact_paper/.

## 2 Results

### 2.1 Methods overview

We first provide an overview of our proposed analytical procedures. The overall workflow of INTACT and INTACT-GSE is summarized in Figure 1. Consider the following structural equation model (SEM), which connects genetic variants (eQTLs and GWAS hits), gene expression of a target gene, and a complex trait of interest,

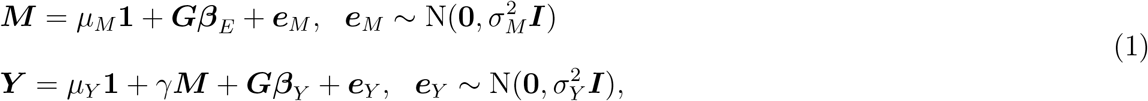

where ***G*** (*n* × *p*), ***M*** (*n* × 1), and ***Y*** (*n* × 1) denote the genotype matrix, the expression levels, and the measurements of a quantitative trait of interest, respectively. The first linear equation is commonly applied in eQTL mapping, and the p-vector, *β_E_*, denotes the true eQTL effects of each genetic variant. The second linear equation assumes that the gene expression levels of the target gene mediate the complex trait with the effect size denoted by *γ*. Additionally, the ***G**β_Y_* term allows separate pleiotropic genetic effects to act on the complex trait (without impacting the expression levels of the target gene). The SEM can be generalized to the two-sample design, where the expression, complex traits, and the corresponding genotypes are measured from two different sets of individuals (Section 1 of Supplemental Methods).

**Figure 1:**
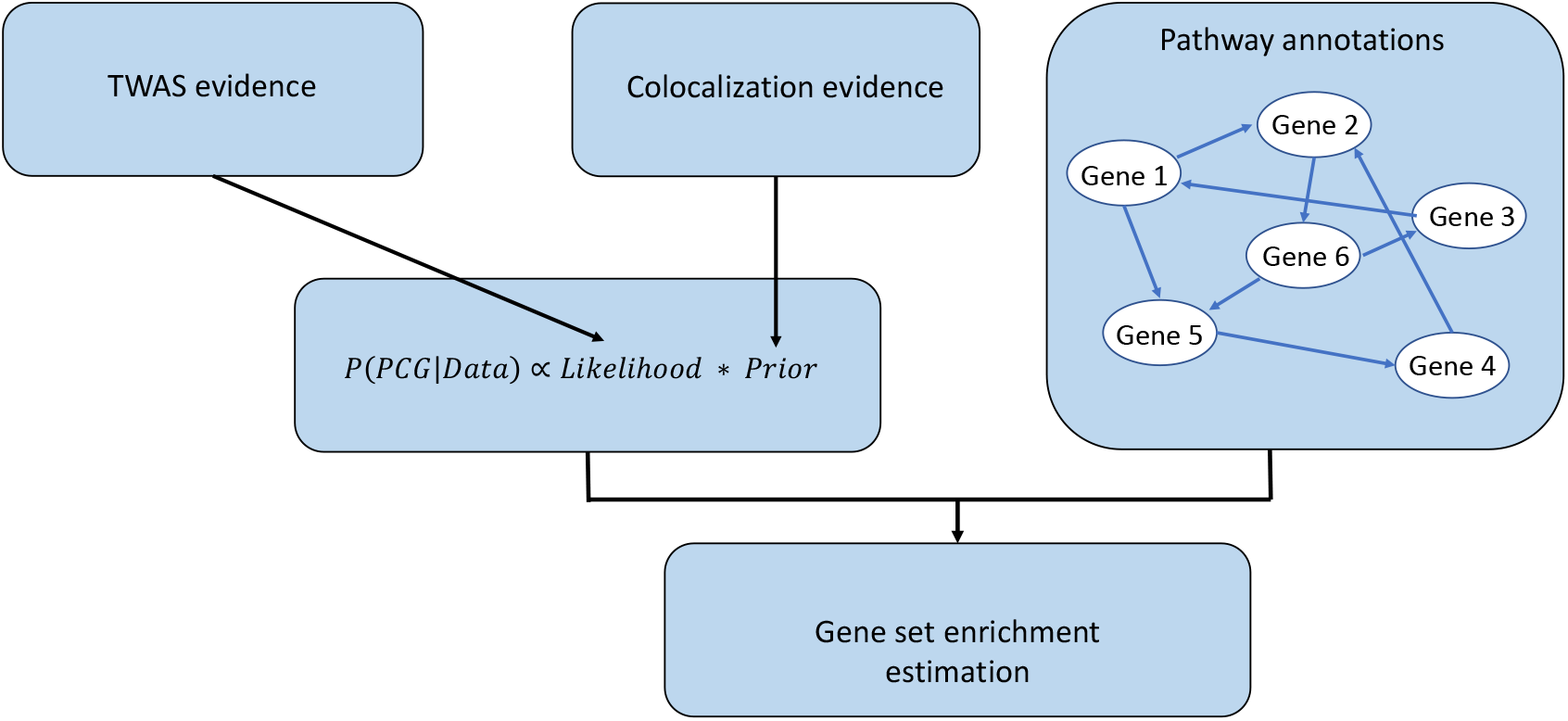
INTACT workflow.

Similar SEMs have been proposed by [14, 9] to implicate PCGs, but with added parametric assumptions to overcome some model identifiability issues (see Methods section). Those additional assumptions can be too stringent to be realistic and become ineffective for handling LD hitchhiking in practice. We avoid these assumptions by making the following observations:

1. The primary goal of the inference is to assess whether *γ* ≠ 0 (but not to estimate *γ*)
2. The SEM (1) implies that the causal eQTLs are colocalized with GWAS hits

The first observation frames PCG implication as a model selection/hypothesis testing problem and distinguishes it from a causal inference problem aiming to estimate γ unbiasedly (e.g., MR-Locus [15]). It is sufficient to establish *γ* ≠ 0 by showing the correlation between the genetically predicted gene expression levels, 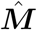, and ***Y*** is non-zero under the necessary assumptions of IV analysis [24, 14, 6]. This procedure has been implemented by various TWAS approaches [7, 8, 23]. Nevertheless, TWAS correlations can occur when *γ* = 0, but causal eQTL SNPs are in LD with causal GWAS SNPs (i.e., in the case of LD hitchhiking; see Proposition 1 of Hukku *et al*. [27] for details). Vanderwheel et al. [24] illustrate that LD hitchhiking leads to the violation of the exclusion restriction (ER) assumption, thus invalidating the putative causality [14, 27].

The key idea of INTACT is to utilize the second observation to constrain the TWAS correlation test results within a formal Bayesian framework. Specifically, we take the TWAS association *z_twas_* score of a target gene and compute a Bayes factor, BF(*z_twas_*) [31], to represent the marginal likelihood of the SEM for *γ* ≠ 0. Furthermore, we formulate an empirical Bayes prior for *γ* ≠ 0 considering the colocalization evidence for the target gene from observed data, i.e.,

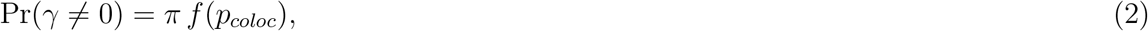

where *π* is an unconstrained TWAS prior (estimated from the data, see the Methods section) and *P_coloc_* represents pre-computed gene-level probabilistic evidence for colocalization of GWAS hits and causal eQTLs. The function, *f* (·), denotes a monotonic mapping such that

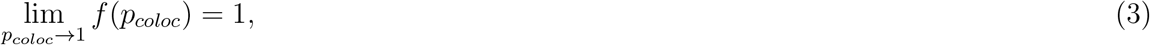

and

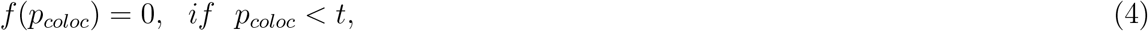

where *t* is a pre-defined non-negative threshold. Equation (4) implements a strong shrinkage effect for genes lacking modest colocalization evidence. Consequently, the posterior probability,

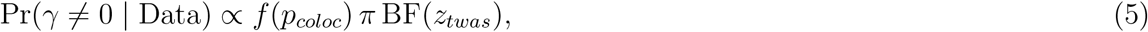

is shrunk to exactly 0 if *P_coloc_* is smaller than the pre-defined threshold *t*, regardless of the likelihood. Because the hitchhiking genes commonly lack colocalization evidence, this analytical strategy effectively removes false-positive findings due to LD-hitchhiking. Here, using a prior specification to implement inference constraints (i.e., the colocalization) is analogous to the Bayesian interpretation of widely-applied shrinkage methods in high-dimensional linear regressions, e.g., ridge and LASSO regressions.

Our software implements a family of *f* (·) functions. Figure 2 provides a visual comparison of the available prior choices. The default prior is the identity mapping truncated at the user-specified threshold, *t* (*t* = 0.05 by default). We refer to the default prior as “linear.” It is worth noting that the inference based on the following step function prior,

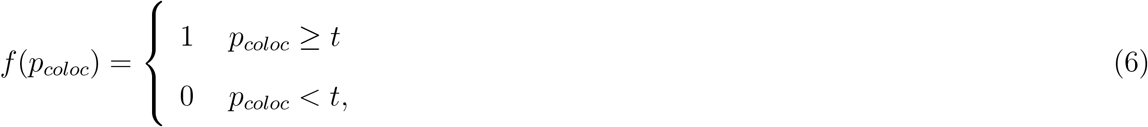

yields comparable results to the *ad hoc* post-processing procedures described in [10, 27] when t is set to 0.50. Additionally, the trivial case in which *f*(*p_coloc_*) ≡ 1 corresponds to a standard Bayesian TWAS analysis.

**Figure 2:**
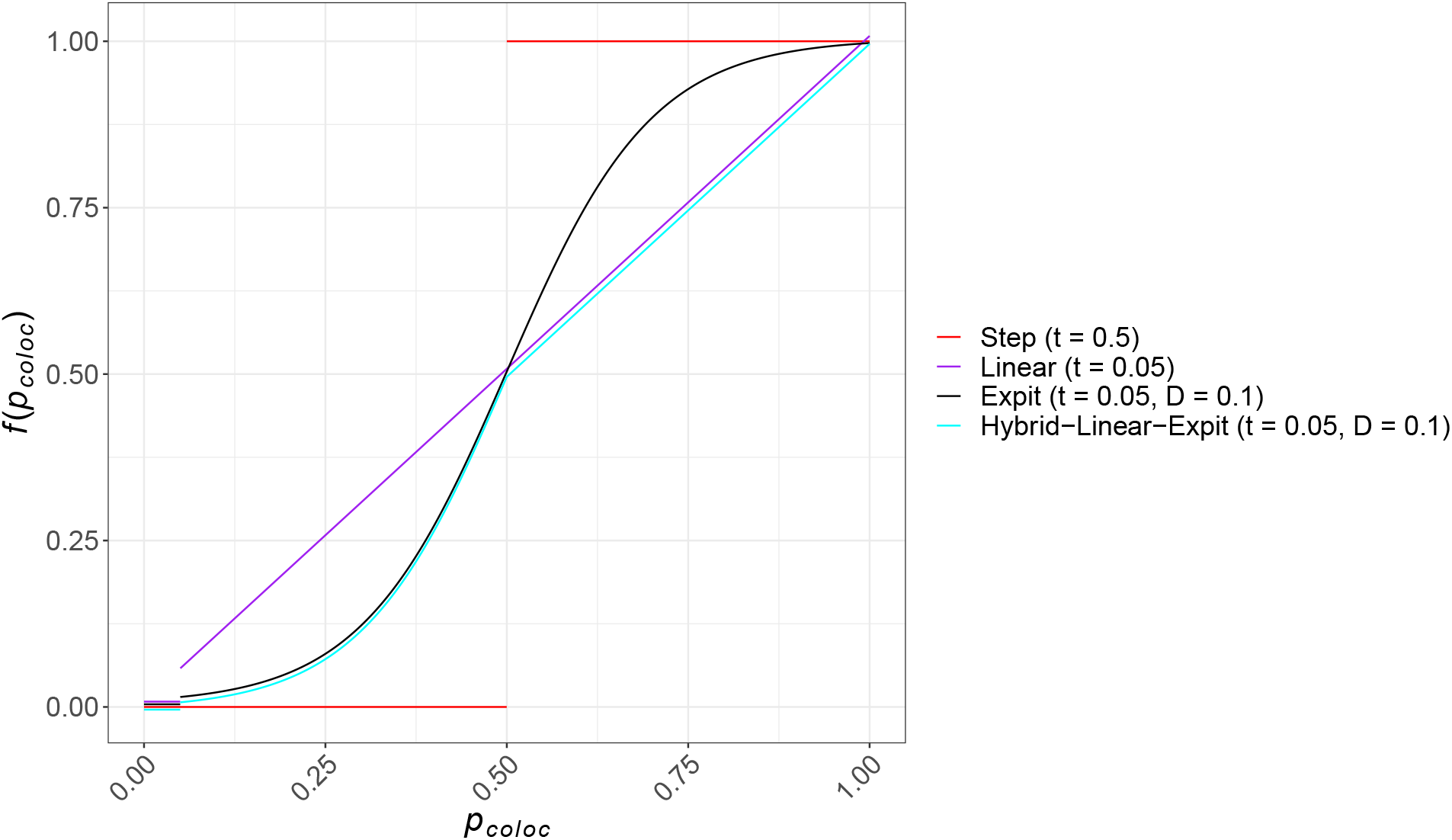
Four prior function options implemented in the INTACT software. Default truncation parameter values *t* and curvature parameter values *D* are included in the legend key, where applicable. (Lines are slightly staggered to prevent overlap for the purpose of visualization.)

### 2.2 Alternative interpretation by Dempster-Shafer theory

We find that the key computational step of integrating colocalization and TWAS evidence in the proposed procedure, Equation (5), has an interesting mathematical connection to the Dempster-Shafer (DS) theory, especially Dempster’s rule of combination. DS theory deals with statistical decision-making utilizing evidence with uncertainty. In our application context, the decision problem can be framed as determining PCGs for a complex trait of interest, and we regard gene-level colocalization and TWAS signals as two sources of relevant but imperfect evidence. In the DS theory, the degree of belief, or mass, for a given type of evidence is represented by a non-negative function ranging from [0, 1]. Hence, we use *m_twas_* and *m_coloc_* to represent the respective mass functions for implicating a gene from TWAS and colocalization analyses. Dempster’s rule of combination states that the overall evidence from multiple sources can be combined by

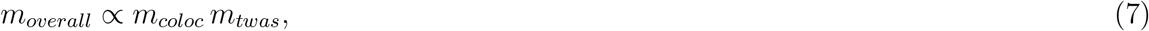

if the two sources of evidence are *DS-independent*, i.e., their *reliability* (in implicating PCGs) is independent [32]. Note that the DS-independence differs from the concept of statistical independence in probability theory (see details in the Methods section).

The unconstrained TWAS posterior probability, *p_twas_*, follows

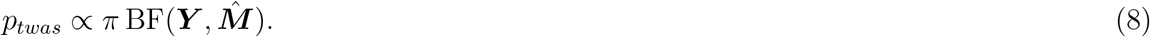

It is rather natural to consider the marginal mass functions

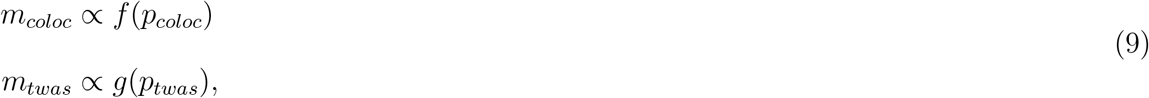

where both *f*(·) and *g*(·) are monotonic transformations of respective posterior probabilities. Importantly, recent work by Hukku *et al*. systematically studies the inferential reproducibility of PCGs implicated by evidence from TWAS scanning and colocalization analysis [27]. Their results lead to the key argument that TWAS and colocalization results are approximately DS-independent. Therefore, within the DS-theory framework, we arrive at the following conclusion,

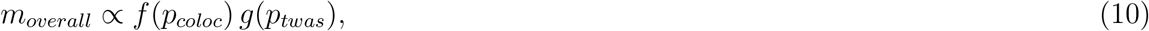

which can be viewed as a special case of Equation 5 (by taking *g*(·) as the identity mapping). This result implies that the rankings of PCG evidence are consistent between the model-based SEM and the non-model-based DS-theory methods for any given datasets.

### 2.3 Probabilistic gene set enrichment analysis

The probabilistic quantification of PCGs enables rigorous gene set enrichment analysis accounting for implication uncertainty. To this end, we propose an expectation-maximization (EM) algorithm, INTACT-GSE, which extends the statistical framework of Bayesian gene set enrichment analysis (BAGSE) [28] to quantify the over-representation of PCGs in a pre-defined gene set.

The key idea of INTACT-GSE is to treat the putative causal status of a given gene as a latent binary indicator. If the indicators are observed, the enrichment analysis of a given gene set can be formulated as a simple estimation problem from a 2 × 2 contingency table. We formulate an EM algorithm to perform maximum likelihood estimation of the corresponding enrichment parameters by treating member genes’ binary putative causal status as missing. We implement multiple numerical routines to quantify the standard errors of the estimated enrichment parameters, including bootstrapping and Fisher scoring methods.

### 2.4 Simulation study

We use simulations to evaluate the performance of INTACT and INTACT-GSE. Our simulated data sets use real genotypes from chromosome 5 of 500 individuals in the GTEx project. The selected genomic region contains 1198 consecutive protein-coding and lincRNA genes with at least 1500 common (MAF > 0.01) cis-SNPs each. We simulate gene expression and complex traits using a scheme similar to [14] and set the signal-to-noise ratios of true genetic associations to match those observed in real data sets. (The details of the simulation procedure are provided in the Methods section.)

#### 2.4.1 Evaluating FDR control and power for implicating genes

We first compare INTACT’s performance as a method for inferring PCGs to some existing computational approaches. The methods for comparison roughly fall into three categories: colocalization-focused approaches represented by fastENLOC [10, 27] with two types of gene-level quantification (GLCP and GRCP); TWAS-only approaches represented by PTWAS and an implementation of LDA MR-Egger/FOCUS algorithm [14, 9]; and the *post hoc* joint TWAS and colocalization analysis method described in Hukku *et al*. [27, 10]. For each dataset consisting of 1198 genes, we set the target FDR control level at 5% for all methods. We then examine the realized FDRs (i.e., 1 – precision) and power (i.e., recall). The results are summarized in Table 1. We observe that TWAS-only approaches suffer excessive false-positive errors due to LD hitchhiking effects. Colocalization-focused methods, on the other hand, have the proper FDR control but are overly conservative with limited power. Both observations are consistent with the conclusions of [27]. The intuitive *post hoc* integration approach yields similar results as INTACT using the step function prior. INTACT using the default prior achieves the best power (~ 40%) among all methods with proper FDR control. Because our simulated data replicate some key characteristics of real data, the estimated power may reflect the practical limitations of the integrative analysis.

**Table 1:**
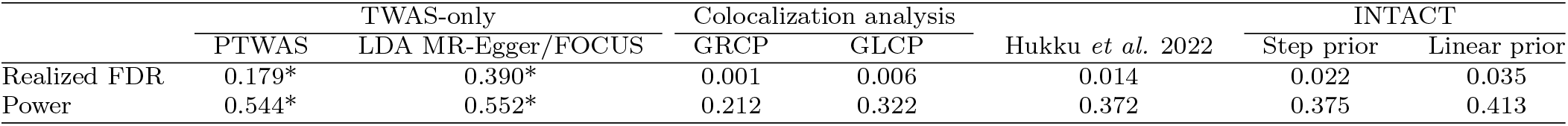
Power and realized FDR for various methods in simulation studies. All methods, except for the *post hoc* procedure by Hukku *et al.,* use the target FDR control level at 5%. The realized FDRs exceeding the target control levels are annotated with asterisks. The Hukku *et al*. procedure intersects genes with GLCP ≥ 0.50 and PTWAS genes rejected at 5% FDR level. INTACT analyses use the step prior (*t* = 0.50) and the default linear prior.

|  | TWAS-only |  | Colocalization analysis |  | Hukku *et al.* 2022 | INTACT |  |
| --- | --- | --- | --- | --- | --- | --- | --- |
|  | PTWAS | LDA MR-Egger/FOCUS | GRCP | GLCP |  | Step prior | Linear prior |
| Realized FDR | 0.179* | 0.390* | 0.001 | 0.006 | 0.014 | 0.022 | 0.035 |
| Power | 0.544* | 0.552* | 0.212 | 0.322 | 0.372 | 0.375 | 0.413 |

We further examine the performance of alternative INTACT prior formulations shown in Figure 2. In our simulation setting, each maintains the proper FDRs and similar power (Supplemental Table S1).

To illustrate the flexibility of INTACT, we alter the TWAS methods for generating *z*-scores. We demonstrate that INTACT is compatible with output from PTWAS, SMR [8], and the LDA MR-Egger/FOCUS model (see Section 2 of Supplemental Methods for our implementation of the LDA MR-Egger/FOCUS model). The resulting FDR and power across different TWAS methods are very similar (Supplemental Table S2).

#### 2.4.2 Gene set enrichment analysis

To evaluate INTACT-GSE’s performance, we randomly annotate 40% of 1,198 genes in each simulated dataset. In our simulation scheme, an annotated gene has a causal effect on the complex trait with probability, 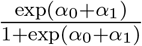, whereas the probability for an unannotated gene is 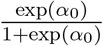. Our focus for this numerical experiment is inference on the enrichment parameter, α_1_, which we select from the set {*α*_1_: 0, 0.25, 0.5, 1.0, 1.5} for each simulated data set. We compare INTACT-GSE with an existing two-stage method in the field [33, 34]. This intuitive approach first classifies each candidate gene based on a 5% FDR threshold using PTWAS p-values; then, it computes an enrichment estimate by constructing a 2 × 2 contingency table, which regards both the binary causal classification and annotation status of each gene as observed.

The point estimates and the corresponding variability from INTACT-GSE and the two-stage algorithms are summarized in Figure 3 and Supplemental Figure S2. For *α*_1_ = 0, both methods yield approximately unbiased estimates. In other scenarios, when *α*_1_ ≠ 0, INTACT-GSE is consistently more accurate, although the estimates from both approaches show downward bias. We attribute the downward bias of INTACT-GSE to its explicit shrinkage step (for stabilizing the estimates) and the imperfect calibration of the input posterior probabilities, which are overly conservative judging by the FDR and power results. Another noticeable advantage of INTACT-GSE over the two-stage approach is its error quantification for estimated *α*_1_ values. By preserving the uncertainty from the classification of PCGs, INTACT-GSE yields more accurate standard error estimates. Consequently, across all enrichment parameter values in our simulation study, the INTACT-GSE yields more calibrated 95% confidence interval with better coverage probabilities than the two-stage approach (Supplemental Table S3).

**Figure 3:**
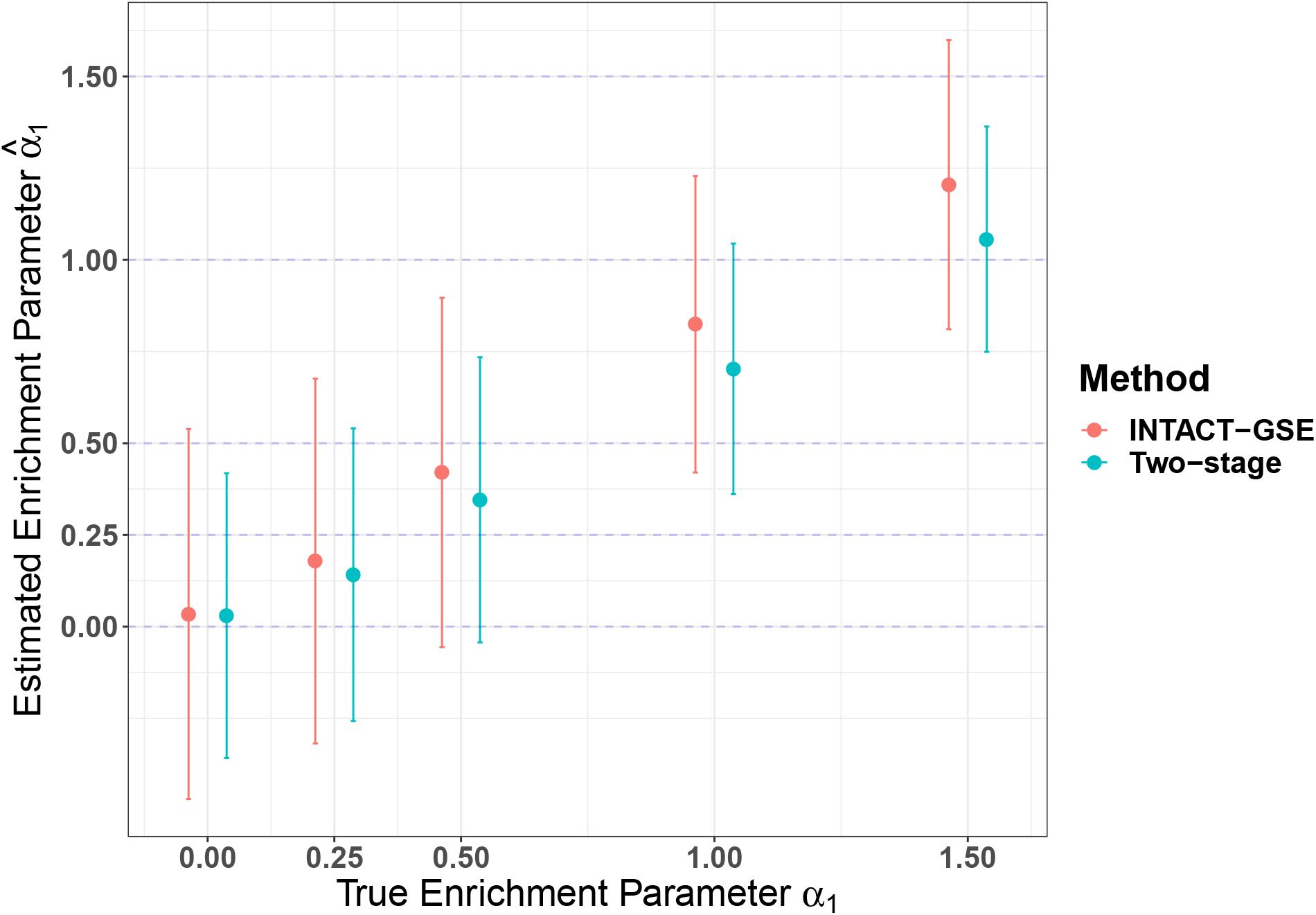
Estimates of gene set enrichment parameter in simulation studies. The estimates and the corresponding 95% confidence intervals from INTACT-GSE and the naive two-stage approach are plotted for each experimented *α*_1_ value. INTACT-GSE yields the most accurate estimates, especially when the true enrichment level is high.

While our primary aim for gene set enrichment analysis is to quantitatively compare and rank different gene sets and pathways via the *α*_1_ estimates, INTACT-GSE results can be trivially applied to test the null hypothesis, *H*_0_ : *α*_1_ = 0. For illustration, we examine the performance of the enrichment testing by INTACT-GSE, the two-stage procedure, and the popular GSEA method [35] (Supplemental Figure S3). We use the unprocessed TWAS *z*-scores as the input for GSEA and evaluate its type I error control. We observe noticeable inflation of the type I error rate to ~ 10% at the 5% control level by the GSEA method (for both weighted and unweighted versions), which is likely due to the LD-hitchhiking effect. In comparison, the two-stage method and INTACT-GSE have proper type I error control. We suspect that in the two-stage method, the effect of excessive false-positive classification errors due to LD-hitchhiking effects is balanced out by the effects of false-negative errors due to the imperfect TWAS analysis power. (We observe that the overall misclassification of genes, including both the type I and type II errors, influences the accuracy of the two-stage gene set enrichment estimation.)

### 2.5 Analysis of complex trait GWAS data from Hukku *et al*. 2022

We apply INTACT to re-analyze the GWAS data from 4 complex traits (HDL and LDL from the GLGC consortium; standing height from the UK Biobank; coronary artery disease from the CARDioGRAM consortium) and the eQTL data across 49 tissues from the GTEx project (v8). Hukku *et al*. [27] report the integrative analysis results of this data set using the *post hoc* approach. The comparison between INTACT and the original Hukku *et al*. analysis is summarized in Supplemental Table S4. Across the 196 tissue-trait pairs, again, we find that INTACT using the step function prior (*t* = 0.50) yields very similar results to the *post hoc* procedure: 98.1% of the findings reported by Hukku *et al*. are also reported by INTACT at the 5% FDR level. At the same FDR control level, INTACT using the default linear prior reports substantially more PCGs. We use this set of INTACT results for the subsequent analysis.

Focusing on coronary artery disease (CAD), we validate the list of genes implicated by INTACT using the DisGeNET database [36], from which a set of 65 genes relevant to CAD with high confidence (DisGenNET scores > 0.3) is curated. INTACT identifies 23 PCGs from this set. The same 23 genes are also implicated by the *post hoc* method and INTACT with the conservative step function prior. Even though there is no qualitative difference in this simple validation by different approaches, the probabilistic quantification of the evidence generated by INTACT is uniquely advantageous for explicit FDR control and subsequent gene set enrichment analysis. To illustrate, we utilize the Comparative Toxicogenomics Database (CTD) [37], which provides curated information about gene-disease relationships integrated with functional and pathway data. CTD ranks pathways according to the available experimental evidence of the pathway’s relevance for a given disease. We query CTD for CAD-related pathways and perform enrichment estimation for the top 10 entries. Coincidentally, all selected pathways are curated from the Reactome database [38]. We apply INTACT-GSE to evaluate each selected pathway in the liver, subcutaneous adipose, visceral adipose, coronary artery, and skeletal muscle tissues, which have been previously implicated as possible relevant tissues for CAD [39, 40]. The results of the enrichment analysis are summarized in Table 2. In summary, INTACT-GSE identifies statistically significant enrichment (at 5% level) in nine pathway-tissue pairs, covering five of the ten examined pathways. In particular, we find that lipid digestion, mobilization, and transportation pathways show strong enrichment in adipose and skeletal muscle tissues. Additionally, the metabolism pathways are enriched in at least one of the five selected tissues. INTACT-GSE does not identify statistically significant enrichment in any tested immune system pathways, possibly due to insufficient power or the relevance of measured transcriptome in CAD etiology.

**Table 2:**
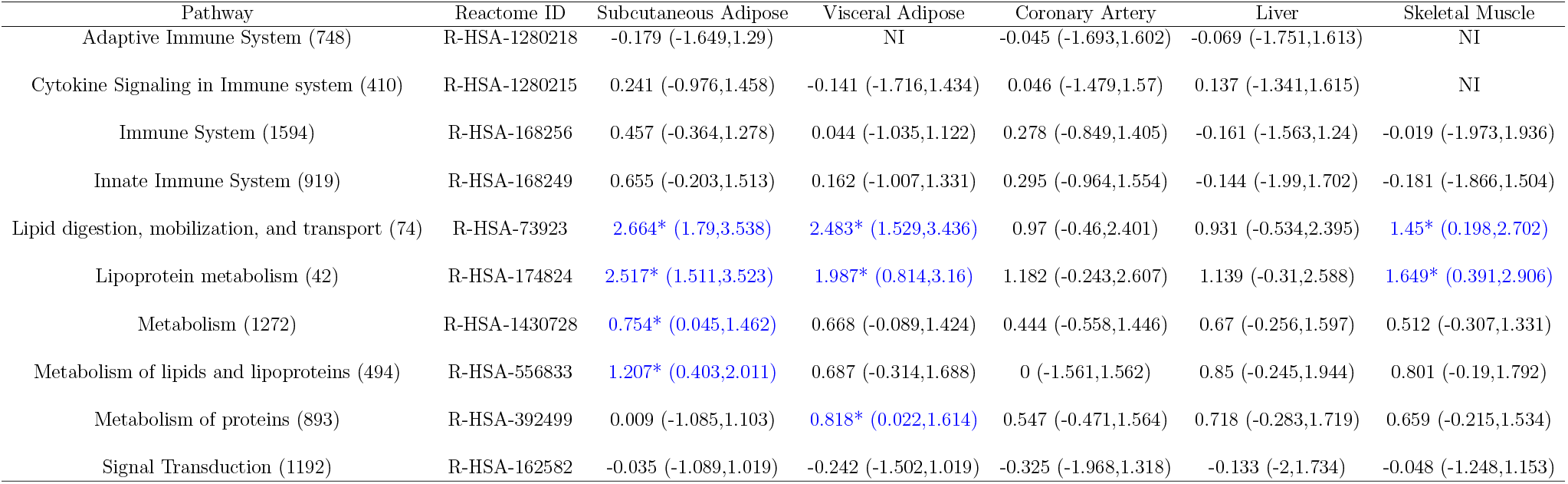
Gene set enrichment estimation for top Reactome pathways relevant to CAD based on CTD. Entries denote the INTACT-GSE estimate of the enrichment level and the corresponding 95% confidence interval. Gene set size (number of ensembl IDs annotated) is shown in the parenthesis beside each pathway name. Statistically significant estimates (at the nominal 5% level) are denoted with an asterisk and highlighted in blue. The estimation results are labeled as Non-informative (NI) if the results are numerically identical to the priors, indicating that likelihood functions from the observed data lack information.

Furthermore, we perform INTACT-GSE analysis for all applicable biological processes Gene Ontology (GO) terms in 196 tissue-trait pairs. The full results are available in Supplemental Excel File 1. In brief, we examine a total of 510714 gene set-tissue-trait combinations, 7333 (or 1.4%) of which show statistically significant enrichment at the nominal 5% level (Supplemental Table S5). Our top enrichment results seemingly reflect some well-documented biological mechanisms underlying CAD. For example, mast cell degranulation processes (GO:0043304) show some of the strongest enrichment levels observed in multiple tissues, and mast cells are known to modulate CAD through multiple pathways in the literature [41, 42].

### 2.6 Analysis of molecular trait data from Sinnott-Armstrong *et al*. 2021

We apply INTACT to the GWAS data of four molecular traits: serum urate, IGF-1, and testosterone levels analyzed in [30]. These complex traits are of interest because their regulating genes and biological pathways are relatively well understood in biochemistry. Sinnott-Armstrong *et al*. interpret GWAS hits and implicate PCGs based on genic distances and existing pathway information. In contrast, we explore the extent to which we can identify relevant genes and pathway interactions by explicitly linking genes to the transcriptome using INTACT. We integrate the GWAS results (i.e., the single-SNP association summary statistics from UK Biobank) for each trait with multi-tissue (49) expression data from GTEx, generating locus-level colocalization and TWAS scan results for each tissue-trait pair. Because the testosterone GWAS analyses are performed separately for males and females, we consider 49 × 4 = 196 tissue-trait pairs in our analysis. We apply INTACT to assess PCGs in each tissue-trait pair using the default linear and the step function priors.

The number of implicated genes at a 5% FDR threshold is included in Supplemental Table S6. In summary, INTACT using the step function prior and Hukku *et al*.’s procedure again yields very similar findings. The INTACT using the default prior implicates substantially more genes, re-confirming our main conclusions from the simulation studies.

We compare the implicated PCGs by INTACT (at the 5% FDR level) and the core genes inferred by the proximity-plus-knowledge-based method using the GWAS data. Core genes, defined in the omnigenic model, are member genes in key biological pathways for relevant traits [43, 30]. Table 3 summarizes the number of genome-wide significant GWAS loci, the uniquely implicated PCGs (in at least one GTEx tissue), and core genes linked to GWAS loci for the four traits. We note that a substantial proportion (in some cases, more than half) of GWAS loci remain unannotated by either approach. The Sinnott-Armstrong *et al*. proximity-plus-knowledge-based approach is limited by the available knowledge of molecular biology underlying each trait (e.g., potentially important yet unknown key pathways). INTACT is constrained by the statistical power of the available molecular QTL data. It is also plausible that gene expression plays insignificant roles in some relevant molecular mechanisms underlying the complex traits of interest (i.e., gene expression levels are not the relevant molecular phenotypes for *all* GWAS loci). Nevertheless, Table 3 shows that INTACT links many more genome-wide significant GWAS loci to candidate genes with strong statistical evidence in TWAS and colocalization analyses. The majority of the PCGs implicated only by INTACT are not considered core genes because they are not included in the pre-defined pathways by Sinnott-Armstrong *et al*. These may be peripheral genes according to the omnigenic model.

**Table 3:** Comparison of significant GWAS loci, implicated PCGs, and linked core genes in Sinnott-Armstrong *et al*. data. The table shows the number of independent and genome-wide significant GWAS loci, the number of unique PCGs implicated by INTACT (at the 5% FDR level and using the default linear prior), and the number of unique core genes annotated in [30] for each trait, respectively. Multiple INTACT PCGs or core genes may be annotated to a GWAS locus.

| Trait | GWAS loci | PCGs by INTACT | Annotated core genes by Sinnott-Armstrong *et al.* |
| --- | --- | --- | --- |
| Serum urate | 299 | 144 | 40 |
| IGF-1 | 402 | 237 | 59 |
| Testosterone (male) | 130 | 69 | 17 |
| Testosterone (female) | 79 | 44 | 8 |

Next, we inspect the overlapping of implicated PCGs by INTACT and the core genes linked to the GWAS signals by Sinnott-Armstrong *et al*. within the pre-defined key pathways. Specifically, we focus on nine sub-pathways relating to the four traits examined by Sinnott-Armstrong *et al*. Because INTACT neither utilizes the known pathway information nor is constrained by a pre-defined genic distance threshold, the implicated PCGs typically represent a different set of core genes in each sub-pathway. Table 4 compares the implicated core genes between INTACT and the proximity-plus-knowledge-based method using the same GWAS summary statistics within the key pathways. Across all traits, INTACT implicates fewer core genes, but in some pathways, the numbers are reasonably close (e.g., the IGF-1 Ras signaling pathway). In addition to the potential relevance of transcriptome data, the type I error control procedure embedded in INTACT may also partially explain the difference (as there is no explicit false discovery control in Sinnott-Armstrong *et al*.‘s method). In many cases, we also observe that the set of PCGs in the examined sub-pathways completely overlaps with the set implicated by the proximity-plus-knowledge-based method. Nevertheless, there are also instances (e.g., Urate Purine metabolism pathway and IGF-1 Ras signaling pathway) in which INTACT and the proximity-plus-knowledge-based method implicate a substantial number of complementary core genes. The detailed breakdown of core genes implicated by each approach is included in Supplemental Excel File 2.

**Table 4:** Comparison of PCGs implicated by INTACT and core genes inferred by the proximity-plus-knowledge-based approach in key pathways. The number of core genes implicated by INTACT represent the implicated PCGs (at the 5% FDR level in at least one GTEx tissue) in each selected key biological pathway. Numbers in parentheses beside pathway names denote the total number of genes annotated for the corresponding pathway in [30].

| Trait | Pathway | Core genes implicated by Sinnott-Armstrong *et al.* 2021 | Core genes implicated by INTACT | Overlap of core genes |
| --- | --- | --- | --- | --- |
| Serum urate | Purine metabolism (88) | 16 | 10 | 4 |
| Serum urate | Solute transport (10) | 8 | 4 | 4 |
| IGF-1 | Growth hormone secretion (32) | 14 | 4 | 4 |
| IGF-1 | IGF-1 secretion (14) | 10 | 2 | 2 |
| IGF-1 | IGF-1 serum balance (18) | 10 | 6 | 6 |
| IGF-1 | Downstream signaling (31) | 9 | 7 | 4 |
| IGF-1 | Ras signaling (156) | 17 | 16 | 6 |
| Testosterone (male) | Synthesis and metabolism (61) | 21 | 7 | 5 |
| Testosterone (female) | Synthesis and metabolism (61) | 13 | 7 | 6 |

Lastly, we apply INTACT-GSE to estimate gene set enrichment for the key sub-pathways of the four traits by Sinnott-Armstrong *et al*. We follow the discussions of tissue relevance in [30] and present the INTACT-GSE results based on the tissue-specific eQTL data for this analysis. Particularly, we use kidney cortex tissue for serum urate, pituitary gland, and liver tissues for IGF-1 (as these tissues are involved in the growth hormone-IGF axis). For testosterone levels, we present the results for tissues relevant to the hypothalamic-pituitary-gonadal (HPG) axis and testosterone breakdown. The enrichment estimation results by INTACT-GSE are summarized in Table 5. Supplemental Excel File 3 provides the complete gene set enrichment results for all 49 GTEx tissues. In 39 pathway-tissue pairs shown in Table 5, the enrichment estimates for 9 pairs are considered statistically significant at the 5% level. For nearly half (17) of the pairs, INTACT-GSE’s estimates are numerically identical to the prior information, indicating that the likelihood surface of the enrichment parameter is flat. Thus, we deem these pathways’ evidence from the observed data non-informative (NI) and label them accordingly. The remaining 13 tested pathways’ estimates are mostly positive but with relatively large standard errors.

**Table 5:** Gene set enrichment estimation of the Sinnott-Armstrong *et al*. data for key biological pathways in relevant tissues. NI indicates that observed data are non-informative for gene set enrichment. Estimates that are significant at the 5% level are highlighted in blue and denoted with an asterisk.

| Trait | Pathway | Tissue |  |  |  |  |
| --- | --- | --- | --- | --- | --- | --- |
| Serum urate | Solute transport (10) | Kidney cortex<br>1.507 (-0.011,3.027) |  |  |  |  |
|  | Purine metabolism (88) | NI |  |  |  |  |
| IGF-1 | Growth hormone secretion (32) | Pituitary gland<br>NI |  | Liver<br>NI |  |  |
|  | IGF-1 secretion (14) | 1.097 (-0.385,2.579) |  | NI |  |  |
|  | IGF-1 serum balance (18) | NI |  | 0.989 (-0.473,2.451) |  |  |
|  | Downstream signaling (31) | 1.229* (0.033,2.426) |  | NI |  |  |
|  | Ras signaling (156) | 0.945* (0.139,1.749) |  | -0.121 (-1.741,1.498) |  |  |
| Testosterone<br>(male) | Synthesis and metabolism (61) | Hypothalamus<br>0.905 (-0.567,2.378) | Pituitary gland<br>0.849 (-0.556,2.255) | Testis<br>0.934 (-0.471,2.340) | Liver<br>1.186 (-0.049,2.423) |  |
|  | Serum homeostasis (5) | 1.667* (0.065,3.269) | 1.596* (0.076,3.115) | NI | NI |  |
|  | HPG signaling (16) | 1.428 (-0.007,2.862) | NI | NI | NI |  |
| Testosterone<br>(female) | Synthesis and metabolism (61) | Hypothalamus<br>NI | Pituitary gland<br>NI | Ovary<br>NI | Adrenal Gland<br>0.932 (-0.473,2.337) | Liver<br>1.194 (-0.211,2.599) |
|  | Serum homeostasis (5) | NI | 1.794* (0.192,3.397) | NI | 1.701* (0.181,3.221) | NI |
|  | HPG signaling (16) | 1.624* (0.195,3.053) | NI | 1.783* (0.306,3.261) | NI | 1.667* (0.197,3.138) |

This analysis highlights some important practical issues in gene set enrichment analysis in such an application context. Particularly, for tissue-specific analysis, comprehensive identification of eQTLs in corresponding tissues is a critical prerequisite for TWAS, colocalization, and down-stream integrative analyses. It is well-known that the power of eQTL discovery is noticeably uneven across human tissues [44] due to the varying difficulty levels in sample collection and tissue biopsy. In our analysis, all the investigated tissues have fewer than half of the complete GTEx samples. For example, the kidney cortex, the key tissue for studying serum urate, has one of the smallest sample sizes (73) among all tissues in the GTEx data. Lack of eQTL discovery leads to uninformative TWAS and colocalization results and subsequently uninformative enrichment analysis results. Consequently, the gene set enrichment analysis should be interpreted cautiously. We emphasize that uninformative results are fundamentally different from the evidence against enrichment. Additionally, protein abundance, rather than transcript abundance, is seemingly a more relevant molecular phenotype for most biochemical interactions represented by the curated pathways. This aspect may also explain reduced statistical power.

## 3 Discussion

This paper introduces a new set of statistical methods to probabilistically integrate TWAS and colocalization results for PCG implications and subsequent gene set enrichment analyses. We demonstrate that our evidence integration procedure, INTACT, identifies genes with higher power than colocalization-focused approaches while properly controlling type I errors commonly induced by LD hitchhiking in TWAS-only methods. The gene set enrichment algorithm, INTACT-GSE, yields more accurate enrichment point estimates with appropriate uncertainty quantification by taking advantage of the probabilistic output from INTACT. It can be used as an important analytical tool to prioritize biological pathways in studying disease etiology.

The INTACT method is motivated by and derived from the SEM (1). Its connection to the Dempster-Shafer theory offers additional justification from a non-generative-model-based perspective and has profound implications for integrating multiple sources of evidence to implicate PCGs. Most importantly, the DS theory, especially Dempster’s rule of combination, does not limit sources of evidence. This feature is most attractive in this context, where multiple relevant molecular phenotypes, such as chromatin structures, DNA methylation levels, and protein abundance, are becoming increasingly available. DS-theory can serve as a foundational statistical framework to simultaneously integrate these types of molecular evidence for gene implication. This future direction is a natural extension of our current work.

Given the technical difficulty and cost of the validation experiments, we prioritize control of false-positive errors in the inference of PCGs. The statistical formulation of PCG discovery is analogous to identifying variant-level genetic associations in traditional statistical genetics. Thus, INTACT emphasizes estimation of false discovery rates from the data via local fdr [45, 46], and our evaluations differ from the treatment of standard binary classification problems in supervised learning, where type I and type II errors are considered indiscriminately (i.e., there is no explicit type I error control).

We consider INTACT a mechanism-aware PCG implication method because the SEM (1) explicitly links the abundance of transcripts to the complex trait of interest via probabilistic modeling. Hence, the scientific interpretation of the inference results is rather natural. Its limitation lies in that transcript abundance is unlikely to be the most relevant molecular phenotype in revealing gene-trait relationships for *all* scenarios. Many authors have shown that isoform usage [47, 48, 49, 50], RNA degradation [51, 52, 53], protein abundance [54, 55, 56], and many other molecular traits [21, 57, 58] play critical roles in the etiology of different diseases. This point is also evidenced by our analyses in which eQTL data only explain a fraction of significant GWAS loci in both Sinnott-Armstrong *et al*. and Hukku *et al*. data. Therefore, applications of INTACT to other available molecular phenotype data may be of interest.

We illustrate that with proper probabilistic quantification of evidence, rigorous gene set enrichment analysis can be performed by applying INTACT-GSE. We note that utilizing the estimated enrichment parameters and gene set information can further boost the discovery of putative causal genes. The idea is to formulate an empirical Bayes prior based on the enrichment estimates and subsequently incorporate it into INTACT. The posteriors for the member genes in an enriched gene set are expected to be increased under these priors. We do not illustrate this feature to make the paper more focused, but similar ideas have been fully demonstrated by [28, 29].

Finally, we acknowledge multiple unresolved practical issues for implicating putative causal genes. First, methodological advancement is unlikely to overcome the lack of quality in genetic association data of molecular and complex traits. As demonstrated through realistic simulations in [26], available genetic association data have poor power for uncovering colocalization instances. Because INTACT relies on colocalization results, it is also limited by this factor. Second, we do not address how to identify trait-relevant tissues (or cell types) in our applications of GTEx data. For INTACT analysis, we utilize available eQTL data across all tissues; for INTACT-GSE analysis, we rely on prior knowledge for tissue specification. Finally, we primarily focus on assessing the presence (or absence) of putative causal links between gene expressions and complex traits, but we do not attempt to quantify the potential causal effect sizes. We acknowledge that causal effect estimation, as in Mendelian randomization (MR) analysis, is an important scientific question. We are committed to addressing this problem in our future work.

## 4 Methods

### 4.1 INTACT method

The complete expression of Equation (5) is given by

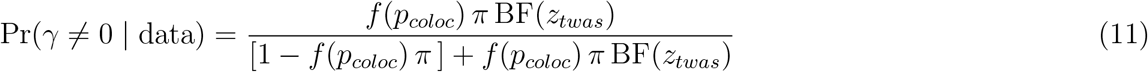

Recall that *f*(*p_coloc_*) represents a monotonic transformation of a gene-level colocalization probability, *p_coloc_*, which quantifies the plausibility of a target gene harboring at least one colocalized variant. Available computational methods, including fastENLOC [10], coloc [12], and eCAVIAR [13], can generate *p_coloc_* results. In the analysis presented in this paper, we use the gene locus-level colocalization probabilities (GLCPs) from fastENLOC as input for INTACT, unless otherwise stated. GLCPs have the advantage of improving the power of colocalization analysis by accommodating the inaccuracy of pinpointing causal variants from genetic association analysis [27]. A valid monotonic mapping for INTACT, *f*, should satisfy properties (3) and (4). Supplemental Figure S1 shows the collection of f functions implemented in our R package, and Section 3 of Supplemental Methods discusses their mathematical properties and considerations for practical usage.

The quantity *π* represents the unconstrained TWAS gene prior. In our implementation, we obtain its lower-bound estimate, 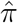, from the observed TWAS *p* values by

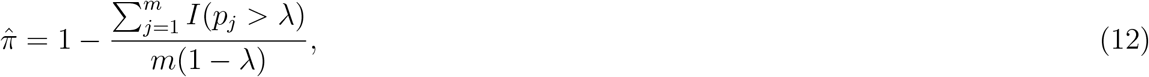

where *m* denotes the total number of genes tested in TWAS, *p_j_* represents the TWAS *p* value for the *j*th gene, and λ is a pre-defined threshold ∈ (0,1). For a better trade-off between the estimation stability and the tightness of the bound, we choose λ = 0.5 by default. This specific estimator is derived from the law of large numbers (LLN) with strong statistical guarantees. It has been made popular by the FDR control method *q*-value [59] and is widely applied in many genomic applications [44, 60].

Finally, we compute the marginal likelihood/Bayes factor for a target gene, BF(*z_twas_*), using the summary association test statistic, *z_twas_*, from TWAS analysis via the Wakefield’s formula [31],

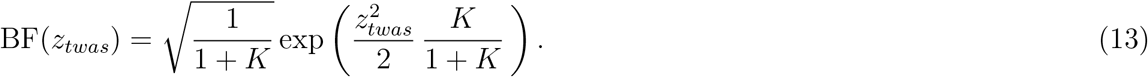

Specifically, hyperparameter *K* represents the prior quantification of association strength. Under the null scenario of no associations, *K* = 0. More generally, 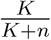 represents the prior expected proportion of variation explained (PVE) by associations, where n is the GWAS sample size. Our software implementation performs Bayesian model averaging over the set {*K* : 1, 2, 4, 8, 16} to cover a spectrum of comprehensive non-null scenarios.

In summary, the INTACT method only requires pre-computed gene-level colocalization probabilities and TWAS *z* scores (from which corresponding *p* values can be derived) for all investigated genes. There are no specific technical requirements for the particular approaches that generate these quantities. Nevertheless, it should be clear that the quality of input directly and profoundly impacts the quality of output by INTACT.

Finally, we note that INTACT can be trivially adjusted and applied to Bayesian TWAS procedures reporting posterior probabilities for candidate genes, e.g., FOCUS [9]. We provide the details in Section 4 of Supplemental Methods.

#### 4.1.1 Comparison to existing methods

Most existing TWAS approaches can be formulated as special cases of SEM (1) and are directly connected to the MR and IV analysis. Particularly, the exclusion restriction (ER), which assumes no pleiotropic effect from the target gene to the complex trait of interest, is required by most existing methods to interpret the observed TWAS associations as potential causal gene-to-trait relationships. The ER is equivalent to assuming *β_Y_* = 0 in the second equation of (1).

A few available approaches for PCG implication, including LDA MR-Egger [14], FOCUS [9], and PMR-Egger [61], aim to relax the ER by working with the full SEM. Nevertheless, all authors recognize that the resulting unrestricted SEM is not identifiable (e.g., multiple combinations of *β_Y_* and *γ* values can lead to the same observed data likelihood). To resolve this issue, the above methods assume that all SNPs in the target gene have the same constant pleiotropic effect, i.e., *β_Y_* = *β_Y_* 1. This assumption is likely motivated by the InSIDE assumption [62, 14, 63] commonly used in MR to relax the ER; however, taking a much stronger form. Because TWAS analysis typically only considers independent eQTLs in the *cis* region of the target gene, the number of independent instruments is typically insufficient for applying the InSIDE assumption in its native form. Such a strong assumption can be useful in some application scenarios, as illustrated by the aforementioned methods. However, it is relatively ineffective in countering the ER violations caused by LD hitchhiking (See Figure 9 of [24] and the relevant discussions therein). The inflation of type I errors due to LD hitchhiking has been well-documented by [14] and [27], and our simulation studies also illustrate this phenomenon. Section 2 of Supplemental Methods provides more technical discussions on this point.

Finally, we find that the fine-mapping strategy utilized by FOCUS is quite effective in detecting and removing between-gene hitchhiking effects [27]. Nevertheless, when considering a single gene, the FOCUS model degenerates to the LDA MR-Egger model, which indicates it is vulnerable to the within-gene LD hitchhiking. The simple solution to this issue is to combine FOCUS and INTACT methods (Section 4 of Supplemental Methods).

### 4.2 Dempster-Shafer theory perspective

DS theory represents a statistical framework of reasoning and decision-making with uncertainty that generalizes Bayesian reasoning. In the DS framework, probabilistic degree of belief, or mass, is assigned to sets of possible outcomes that comprise a system. In our notations, *m_twas_* and *m_coloc_* represent the degree of belief that a target gene is PCG given its TWAS and colocalization evidence, respectively. Dempster’s rule of combination integrates multiple sources of probabilistic information under the constraint that they are DS-independent; that is, they have independent *reliability* for reaching the same decision (in our case, implicating a PCG) [32]. We reason that TWAS and colocalization evidence is approximately DS-independent because their inferential reproducibility for implicating PCGs is low in practice. Particularly, Hukku et al. show that genes with strong TWAS evidence often show weak colocalization evidence (due to LD hitchhiking); genes with strong colocalization evidence can have weak TWAS evidence due to the cancellation effects of horizontal pleiotropy. Overall, the two sets of PCGs separately implicated by TWAS and colocalization evidence are largely non-overlapping. Therefore, it can be argued that their reliability in implicating PCG is approximately independent, i.e., *m_twas_* and *m_coloc_* are approximately DS-independent. It is also important to distinguish the concepts of statistical and DS independence. Types of evidence can be DS independent but statistical dependent. A commonly cited example of such a scenario is that different eyewitness testimony for the same event are statistically dependent, but they are often regarded as DS-independent in decision-making. Similarly, in our application, TWAS and colocalization evidence are also statistically dependent, as they are often derived from the same dataset.

### 4.3 INTACT-GSE method

Given a pre-defined gene set of interest, we estimate the enrichment of PCGs using an EM algorithm. Specifically for gene *i*, we model the relationship between the latent PCG indicator, *γ_i_*, and the corresponding gene set membership indicator, *d_i_*, by the following logistic prior function,

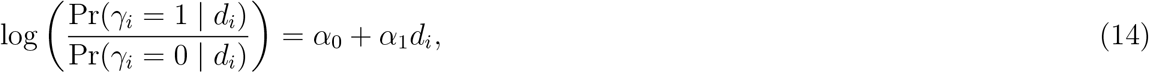

where *α*_1_ represents the log odds-ratio of an annotated gene being a PCG versus an unannotated gene and is of interest for inference.

We design an expectation-maximization (EM) algorithm to find the maximum likelihood estimates for the parameter *α* = (*α*_0_, *α*_1_) by treating *γ_i_*’s as missing data. We provide the full technical details of the EM algorithm in Section 5 of Supplemental Methods. Briefly, the E-step is a simple application of the Bayes rule for computing conditional expectations of *gamma_i_*’s, and the M-step is equivalent to performing inference in a 2 × 2 contingency table, where the cell counts are fractional. The EM algorithm reports the MLE, 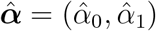, and the corresponding variance. (We denote the variance of 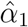 by 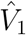.)

In practice, the observed data can be uninformative for the enrichment analysis, which corresponds to the scenario in which the underlying latent 2 × 2 contingency table is highly imbalanced and some cell counts are near 0 (e.g., with few PCGs and/or few annotated genes). In this scenario, the resulting 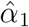 is typically unstable (i.e., extremely slow to converge) and with a large standard error. To accommodate such scenarios, we assign a N(0, 1) prior to *α*_1_. By assuming MLE is approximately normal, we report the resulting posterior mean,

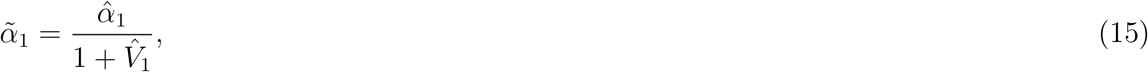

and the corresponding variance,

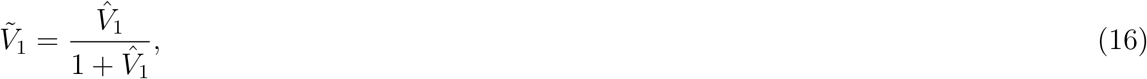

as the result for our enrichment analysis. For less informative data (i.e., *V*_1_ ≫ 1), our enrichment estimate is shrunk back to the prior. Conversely, when the observed data is highly informative, the MLE from the EM algorithm dominates.

### 4.4 Simulation studies

We design simulations to assess the performance of INTACT and INTACT-GSE. For each simulated data set, we use real genotypes of all SNPs on chromosome 5 from 500 individuals in the GTEx (v8) data. The selected genomic region contains 1198 consecutive genes with at least 1500 common (MAF > 0.01) *cis*-SNPs each. For each gene, we randomly select 2 causal eQTLs, *g*_1_ and *g*_2_, and a distinct causal GWAS SNP, *g*_3_. We also generate a gene set annotation for the *i*-th gene by randomly drawing

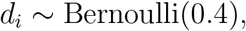

i.e., on average, 40% of the 1198 genes are annotated. Subsequently, the *i*-th gene is labeled as a PCG with the probability

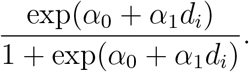

We fix *α*_0_ = −1.735 and vary *α*_1_ across simulations.

Given (*d_i_, γ_i_*) for each gene, we simulate the expression and the complex trait data using the SEM (1). Specifically, all genetic effects (*beta*’s) and non-zero *γ* are drawn from a N(0, *ϕ*^2^) distribution, and the residual error variances are set to 1. We find that setting *ϕ* = 0.6 best matches the simulated data to the observed genetic association data in the distributions of single-SNP *z* scores.

Using this scheme, we generate 100 datasets for each *α*_1_ ∈ {0.00, 0.25, 0.50,1.00,1.50}.

To analyze the simulated data, we perform eQTL and GWAS fine-mapping analysis using DAP-G [25]. The fine-mapping results are subsequently used for TWAS and colocalization analyses using PTWAS and fastENLOC, respectively. We also perform alternative TWAS analysis using SMR and LDA MR-Egger algorithms. To compare gene set enrichment methods, we apply weighted and unweighted GSEA using the R software package fgsea (v1.12.0) with its default setting.

### 4.5 Real data analysis

#### 4.5.1 Processing of Hukku *et al*. data

We download and use TWAS scan and colocalization results from Hukku *et al*. [27] for INTACT analysis. For CAD gene set enrichment analyses, we obtain a list of biological process (BP) GO terms and Reactome pathways with corresponding gene Ensembl ID members using the `biomaRt` R package (v2.42.1). In total, we consider 12,577 BP GO terms. We do not estimate enrichment for all BP GO terms in each tissue. Those with fewer than 70% of genes represented among a data set are not considered. Additionally, we test enrichment for the BP GO terms for the three other complex traits. Again, tissue-trait-GO term combinations for which fewer than 70% of GO term genes are represented among the data are not considered.

#### 4.5.2 Processing of Sinnott-Armstrong *et al*. data

We download several data files provided by Sinnott-Armstrong *et al*. [30] in our analysis. All genomic data are lifted over to GRCh38 to match the GTEx (v8) reference build. To generate colocalization and TWAS-scan results for each of the four molecular GWAS traits, we run fastENLOC and PTWAS using the converted GWAS summary statistics and the GTEx multi-tissue eQTL data.

#### 4.5.3 Analysis of proximal genes for GWAS hits

To generate Table 3, we use Supplementary Files 1-4 from [30], which contain lead SNPs from loci with GWAS hits for each of the four molecular traits. To generate the middle column of Table 3, we first generate a list of gene Ensembl IDs that INTACT implicates in at least one GTEx tissue using a 5% FDR threshold in each tissue-trait pair. We convert the Ensembl IDs to gene symbols using the `biomaRt` R package. Ensembl IDs without a matching gene symbol are dropped. Then, we intersect lead GWAS hit coordinates with gene coordinates for the INTACT hits using the `intersect` function from `bedtools` (v2.22.0) to find the number of lead SNPs proximal to an INTACT gene hit. Gene coordinates are obtained from the file all_hits_msiggenes.bed from [30]. Because we aim to identify GWAS hits *proximal* to implicated genes, we extend gene coordinates by 100 kb on either end (truncating at the chromosome ends). To generate the right-most column of Table 3, we first obtain a list of core genes for each molecular trait from Supplementary Files 8-10 from [30]. We intersect the coordinates of each set of GWAS hits with the corresponding set of extended coordinates of core genes.

#### 4.5.4 Core gene pathway enrichment analysis

We estimate gene set enrichment using the core gene pathway information provided in Supplementary Files 8-10 of [30]. INTACT gene-level probabilistic results for each tissue-trait pair are generated using a linear prior. Gene symbols from the supplementary files are converted to Ensembl IDs before estimating enrichment for each pathway.

