## Supplemental Materials for "Probabilistic integration of transcriptome-wide association studies and colocalization analysis prioritizes molecular pathways of complex traits"

### Supplemental Figures

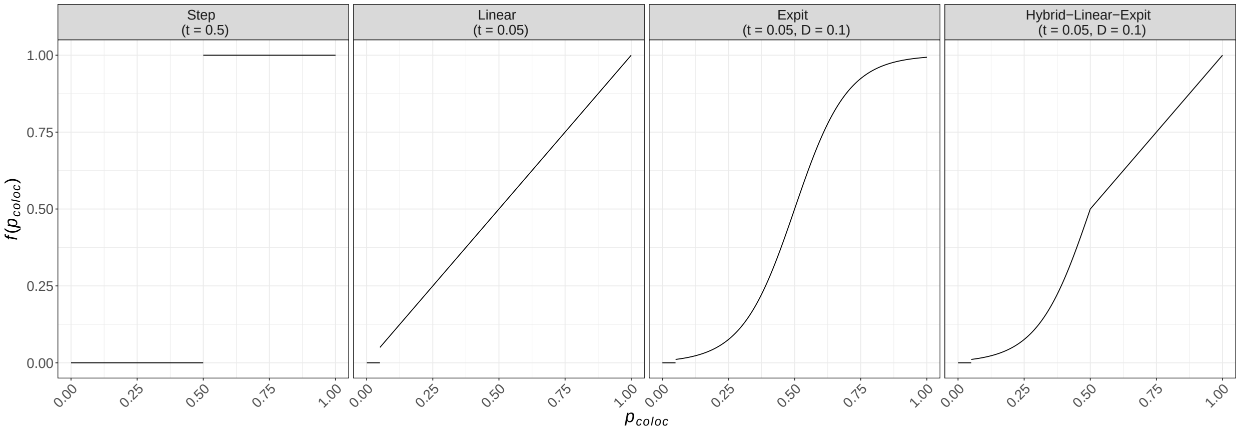


Figure S1: **Family of prior functions included in the INTACT software package** Default truncation parameter $t$ and curvature parameter $D$ values are given in parentheses under each prior name, where applicable.

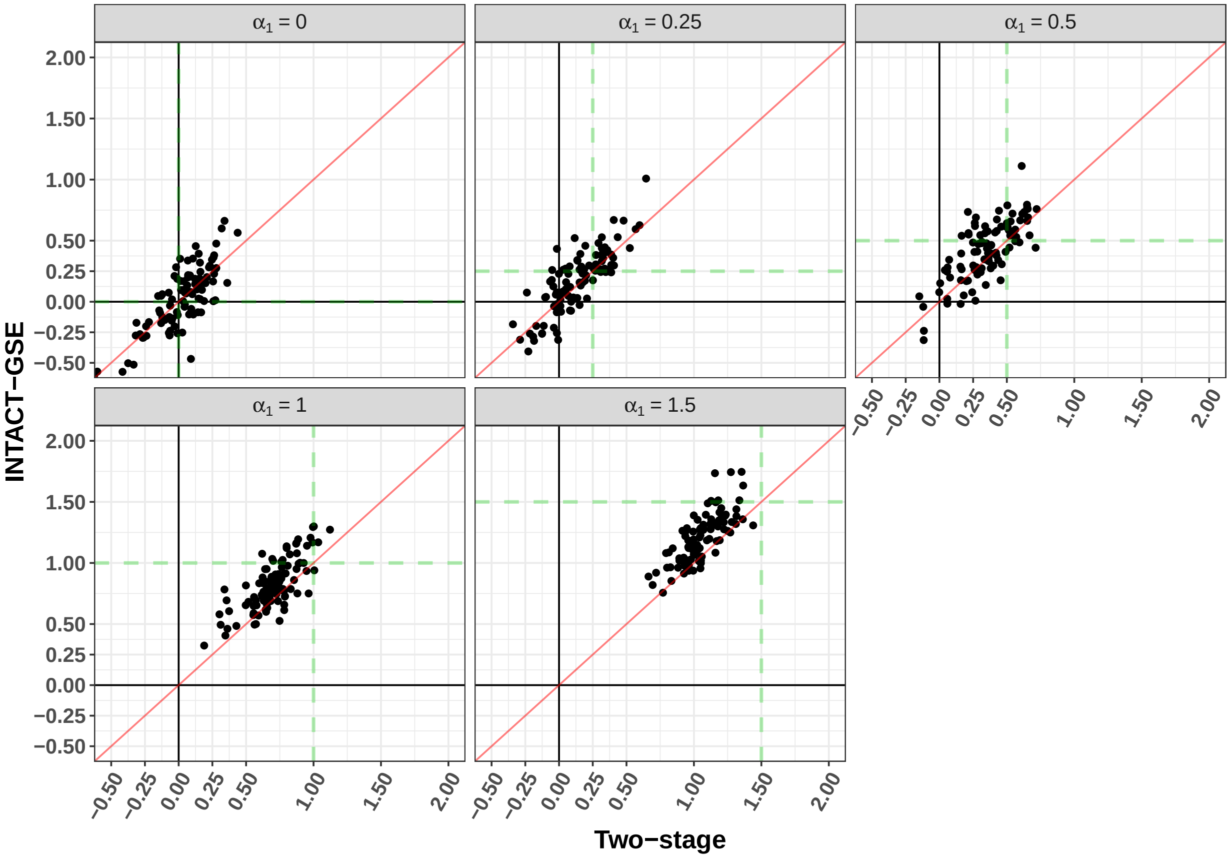


Figure S2: **Comparison of point enrichment estimates by INTACT-GSE and the two-stage approach for each simulated data set** Each point represents the estimate from one simulated data set, and panels correspond to true enrichment parameters. Green dashed lines denote the true $\alpha_1$ values. It is clear that the two-stage method is more conservative when $\alpha_1 > 0$.

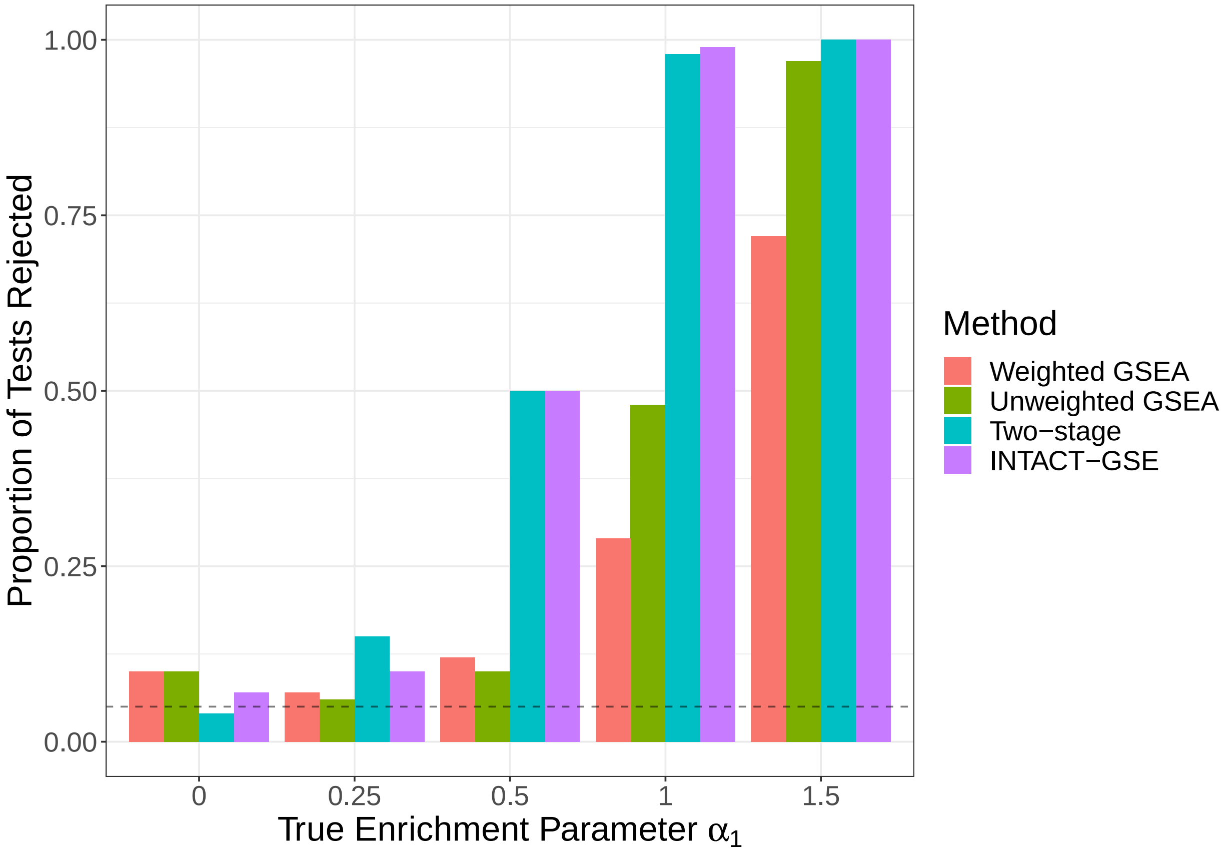


Figure S3: **Comparison of enrichment testing performance by INTACT-GSE, two-stage method, and weighted/unweighted GSEA methods** The y-axis shows the proportion of tests with p-value $< 0.05$.

### Supplemental Tables

Table S1: **Power and realized FDR of INTACT using different prior functions** The step prior is truncated at $t = 0.5$. The linear, expit, and hybrid priors use a truncation threshold of $t = 0.05$. For the expit priors, $D = 0.1$. The target FDR control level is at 5%.

| | Step | Linear | Expit | Hybrid-Linear-Expit |
|---|---|---|---|---|
| Realized FDR | 0.022 | 0.035 | 0.028 | 0.026 |
| Power | 0.375 | 0.413 | 0.417 | 0.416 |

Table S2: **Power and realized FDR of INTACT using z-scores generated from different TWAS methods** Three types of TWAS $z$ scores are generated from PTWAS, SMR, and the LDA-Egger/Focus methods. The target FDR control level is at 5%.

| | PTWAS | SMR | LDA-Egger/FOCUS |
|---|---|---|---|
| Realized FDR | 0.035 | 0.034 | 0.044 |
| Power | 0.413 | 0.41 | 0.421 |

Table S3: **Coverage probability of confidence intervals by enrichment estimation procedures** Each entry represents the coverage probability of the reported 95% confidence intervals over 100 simulations. Standard errors for INTACT-GSE are computed via numerical differentiation of the Fisher score vector and more accurate.

| | True enrichment parameter ($\alpha_1$) value | | | | |
|---|---|---|---|---|---|
| Method | 0 | 0.25 | 0.50 | 1.00 | 1.50 |
| INTACT-GSE | 0.93 | 0.91 | 0.91 | 0.88 | 0.64 |
| Two-stage | 0.96 | 0.89 | 0.82 | 0.53 | 0.18 |

Table S4: **Significant PCG findings of Hukku *et al.* data across 196 trait-tissue pairs** The analysis is performed for 4 complex traits and 49 GTEx tissues. The significant INTACT PCGs are identified at the 5% FDR level in each trait-tissue pair. The Hukku *et al.* method identifies noteworthy genes by intersect significant TWAS genes (at the 5% FDR level) and the genes with gene-level colocalization probability, GLCP, > 0.50. The numbers in parentheses denote the unique genes across 49 tissues.

| | | INTACT | |
|---|---|---|---|
| Trait | Hukku *et al.* 2022 | Step prior ($t = 0.50$) | Default linear Prior |
| Heights | 4723 (966) | 5056 (1043) | 14238 (2425) |
| HDL | 454 (128) | 396 (111) | 988 (204) |
| LDL | 346 (88) | 300 (79) | 649 (146) |
| CAD | 801 (179) | 797 (178) | 1498 (291) |
| Total | 6324 | 6549 | 17373 |

Table S5: **Gene set enrichment analysis results for biological processes GO terms in CAD, HDL, LDL, Heights, and LDL** For each trait, GO terms are tested in each of the 49 GTEx tissues for which the term has more than 70% of genes represented among the tissue-trait-specific TWAS and colocalization summary data. The number of GO terms tested is the total number of GO term-tissue pairs that meets this criteria. Enrichment estimates are deemed significant based on a nominal p-value threshold of 5%.

| Trait | Number of GO terms tested | Number of significant enrichment estimates |
|---|---|---|
| CAD | 280837 | 6393 |
| HDL | 99469 | 603 |
| Heights | 46876 | 21 |
| LDL | 83532 | 316 |
| Total | 510714 | 7333 |

Table S6: **Significant PCG findings of Sinnott-Armstrong *et al.* data across 196 trait-tissue pairs** The significant INTACT PCGs are identified at the 5% FDR level in each trait-tissue pair. The Hukku *et al.* method identifies noteworthy genes by intersecting significant TWAS genes (at the 5% FDR level) and the genes with gene-level colocalization probability, GLCP, > 0.50. The numbers in parentheses denote the unique genes across 49 tissues.

| | | INTACT | |
|---|---|---|---|
| Trait | Hukku *et al.* 2022 | Step prior ($t = 0.50$) | Default linear prior |
| Serum urate | 1522 (444) | 1581 (466) | 8266 (1623) |
| IGF-1 | 2192 (575) | 2289 (599) | 11667 (2033) |
| Testosterone (male) | 1241 (316) | 1297 (321) | 5090 (903) |
| Testosterone (female) | 579 (160) | 551 (154) | 2216 (491) |
| Total | 5534 | 5718 | 27239 |

### Supplemental Methods

### 1 SEM for two-sample design

Consider applying the SEM described in the main text to the two cohorts of non-overlapping individuals. For the first cohort, we obtain

$$\begin{aligned} \boldsymbol{M}_1 &= \mu_{M,1}\mathbf{1} + \boldsymbol{G}_1\boldsymbol{\beta}_E + \boldsymbol{e}_{M,1}, \quad \boldsymbol{e}_{M,1} \sim \mathrm{N}(\mathbf{0}, \sigma^2_{M,1}\boldsymbol{I}) \\ \boldsymbol{Y}_1 &= \mu_{Y,1}\mathbf{1} + \gamma\boldsymbol{M}_1 + \boldsymbol{G}_1\boldsymbol{\beta}_Y + \boldsymbol{e}_{Y,1}, \quad \boldsymbol{e}_{Y,1} \sim \mathrm{N}(\mathbf{0}, \sigma^2_{Y,1}\boldsymbol{I}) \end{aligned} \tag{S1}$$

Similarly, for the second cohort,

$$\begin{aligned} \boldsymbol{M}_2 &= \mu_{M,2}\mathbf{1} + \boldsymbol{G}_2\boldsymbol{\beta}_E + \boldsymbol{e}_{M,2}, \quad \boldsymbol{e}_{M,2} \sim \mathrm{N}(\mathbf{0}, \sigma^2_{M,2}\boldsymbol{I}) \\ \boldsymbol{Y}_2 &= \mu_{Y,2}\mathbf{1} + \gamma\boldsymbol{M}_2 + \boldsymbol{G}_2\boldsymbol{\beta}_Y + \boldsymbol{e}_{Y,2}, \quad \boldsymbol{e}_{Y,2} \sim \mathrm{N}(\mathbf{0}, \sigma^2_{Y,2}\boldsymbol{I}) \end{aligned} \tag{S2}$$

Note that the two sets of SEMs share only parameters of interest, $\boldsymbol{\beta}_E$ and $\gamma$, other parameters, e.g., intercepts and residual error variances, can be sample-specific. The sample sizes in the two cohorts are generally different.

In practice, the two different cohorts are dedicated to studying different phenotypes. For example, the first cohort is typically designed for eQTL mapping and the second cohort is for GWAS. Thus, the observed data is known from a two-sample design, i.e.,

$$\begin{aligned} \boldsymbol{M}_1 &= \mu_{M,1}\mathbf{1} + \boldsymbol{G}_1\boldsymbol{\beta}_E + \boldsymbol{e}_{M,1}, \quad \boldsymbol{e}_{M,1} \sim \mathrm{N}(\mathbf{0}, \sigma^2_{M,1}\boldsymbol{I}) \\ \boldsymbol{Y}_2 &= \mu_{Y,2}\mathbf{1} + \gamma\boldsymbol{M}_2 + \boldsymbol{G}_2\boldsymbol{\beta}_Y + \boldsymbol{e}_{Y,2}, \quad \boldsymbol{e}_{Y,2} \sim \mathrm{N}(\mathbf{0}, \sigma^2_{Y,2}\boldsymbol{I}) \end{aligned} \tag{S3}$$

This is a common scenario in TWAS. Although $\boldsymbol{M}_2$ is unobserved, its predictive value, $\hat{\boldsymbol{M}}_2$, can be computed by learning from the eQTL data in the first cohort. The TWAS association $z$-score, $z_{twas}$, is then computed by using $\boldsymbol{Y}_2$ and $\hat{\boldsymbol{M}}_2$. The key point is that under the two-sample design and SEM (S3), BF($z_{twas}$) remains a valid likelihood function for the shared common parameter $\gamma$.

#### 2 Alternative SEM fitting approach

We implement an alternative SEM fitting algorithm to represent the existing TWAS approaches, namely LDA MR-Egger and FOCUS, allowing additional pleiotropic effects. Many authors [1, 2, 3] note that the original SEM is not identifiable without further parametric assumptions. This is because any combinations of $(\gamma, \boldsymbol{\beta}_Y)$ leading to the same value of $(\gamma\boldsymbol{\beta}_E + \boldsymbol{\beta}_Y)$ result in the same likelihood values. To resolve the identifiability issue, existing methods make an additional assumption that all SNPs have the constant pleiotropic effects, i.e.,

$$\boldsymbol{\beta}_Y = \alpha\mathbf{1}. \tag{S4}$$

With this assumption, the SEM can be straightforwardly fitted, and a unique $\gamma$ can be estimated from the observed data. We implement this algorithm to fit individual-level TWAS data in our simulation studies and represent LDA MR-Egger and FOCUS methods.

The assumption (S4) closely resembles the assumption of burden test in genetic association analysis of rare variants. However, its validity is difficult to confirm from the available GWAS data, especially for common SNPs. More importantly, simple analysis can show that the assumption does not resolve LD hitchhiking. Consider in a true generative model, $\gamma = 0$ and a true GWAS hit with $\beta_Y \neq 0$ is in LD with the causal eQTLs. The SEM with assumption (S4) becomes misspecified. The GWAS effect can only be partially explained by the pleiotropic effect (by regressing $\boldsymbol{Y}$ on $\sum_i^p \boldsymbol{g}_i$), and the $\gamma$ estimate (by regressing $\boldsymbol{Y}$ on $\hat{\boldsymbol{M}}$) is unlikely 0, which leads to false-positive PCG findings.

#### 3 Prior choices for INTACT

Below we list the options for the prior function $f(\cdot)$ included in the software implementation of our method. The parameter $t$ is a truncation threshold. The default $t$ value for the step prior is 0.5, and for all other priors, the default is 0.05. The parameter $D$ controls the steepness of the expit prior curve. As $D$ decreases, colocalization probabilities in the range $(t, 0.5)$ are shrunk closer to zero, resulting in a more-conservative prior. The default $D$ value is 0.1.

**Step**:

$$f(p_{coloc}) = \mathbf{1}(p_{coloc} \geq t) = \begin{cases} 1 & \text{if } p_{coloc} \geq t \\ 0 & \text{otherwise} \end{cases}$$

**Linear**:

$$f(p_{coloc}) = p_{coloc}\,\mathbf{1}(p_{coloc} \geq t)$$

**Expit**:

$$\begin{aligned} f(p_{coloc}) &= \text{expit}\left[(p_{coloc} - 0.5)/D\right]\,\mathbf{1}(p_{coloc} \geq t) \\ &= \frac{1}{1 + \exp\left(-\frac{p_{coloc} - 0.5}{D}\right)}\,\mathbf{1}(p_{coloc} \geq t) \end{aligned}$$

**Hybrid-Linear-Expit**:

$$f(p_{coloc}) = \begin{cases} 0 & p_{coloc} < t \\ \text{expit}\left[(p_{coloc} - 0.5)/D\right] & t \leq p_{coloc} < 0.5 \\ p_{coloc} & p_{coloc} \geq 0.5 \end{cases}$$

All prior functions utilize hard thresholding to shrink the prior to exactly 0 for weak colocalization evidence, a necessity for guarding against LD hitchhiking. They differ in dealing with modest to strong colocalization evidence. noticeably, the step function can be viewed as a special case of the expit function when $D \to \infty$. Comparing to the neutral linear function, the expit family prior up-weight colocalization evidence when $p_{coloc} > 0.5$ and down-weight it when $p_{coloc} < 0.5$. This can be a desired property, provided that colocalization probabilities are known to be overly conservative in true colocalization instances [4]. The hybrid function only down-weights modest colocalization evidence when $p < 0.50$ but remains neutral for strong evidence represents an overly conservative prior choice.

#### 4 INTACT algorithm for Bayesian TWAS result

In the main text, we illustrate the INTACT algorithm assuming output from frequentist TWAS methods (in the forms of $z$ scores or $p$ values). Here we show the INTACT algorithm for Bayesian TWAS methods, which report a posterior probability for each candidate gene (e.g., FOCUS).

Consider the TWAS prior probability, $\pi_i$, is available for gene $i$. The TWAS Bayes factor, $\text{BF}_i$, can be computed by

$$\text{BF}_i = \frac{p_{twas,i}}{1 - p_{twas,i}} \frac{1 - \pi_i}{\pi_i}, \tag{S5}$$

where $p_{twas,i}$ denote the posterior TWAS probability for gene $i$. Given $\pi_i$ and $\text{BF}_i$, INTACT incorporates gene-level colocalization probability, $p_{coloc,i}$, by

$$\Pr(\gamma_i \neq 0 \mid \text{ Data}) \propto \pi_i \, f(p_{coloc,i}) \, \text{BF}_i, \tag{S6}$$

i.e., INTACT modifies the original prior by rescaling it according to the colocalization evidence.

#### 5 INTACT-GSE EM algorithm

Given the INTACT output posteriors, $\{\Pr(\gamma_i \neq 0 \mid \text{ data }) : i = 1, 2, ..., m\}$, for $m$ genes. We first compute a consistent exchangeable prior (with respect to gene set annotations) by the total expectation law and law of large numbers,

$$\Pi = \Pr(\gamma \neq 0) = \text{E}\left[\Pr(\gamma \neq 0 \mid \text{ data })\right] \approx \frac{1}{m} \sum_i \Pr(\gamma_i \neq 0 \mid \text{ data }). \tag{S7}$$

The likelihood evidence for each gene $i$ subsequently can be computed by a Bayes factor, $\text{BF}_i$, i.e.,

$$\text{BF}_i = \frac{\Pr(\gamma_i \neq 0 \mid \text{ data })}{1 - \Pr(\gamma_i \neq 0 \mid \text{ data })} \frac{1 - \Pi}{\Pi} \tag{S8}$$

We initiate the EM algorithm by setting

$$\begin{aligned} \alpha_0^{(0)} &= \log\left[\frac{\sum_{i=1}^m \Pr(\gamma_i \neq 0 \mid \text{ data })}{m - \sum_{i=1}^m \Pr(\gamma_i \neq 0 \mid \text{ data })}\right] \\ \alpha_1^{(0)} &= 0 \end{aligned} \tag{S9}$$

In the E-step, we update the prior for each gene $i$ according to its annotation status, $d_i$, and the current enrichment parameter estimate, $(\alpha_0^{(t)}, \alpha_1^{(t)})$, i.e.,

$$\pi_i^{(t)} = \Pr(\gamma_i = 1 \mid d_i, \alpha_0^{(t)}, \alpha_1^{(t)}) = \frac{\exp(\alpha_0^{(t)} + \alpha_1^{(t)} d_i)}{1 + \exp(\alpha_0^{(t)} + \alpha_1^{(t)} d_i)}. \tag{S10}$$

The updated conditional expectation for the missing data, $\gamma_i$, is updated using Bayes rule,

$$\mathrm{E}(\gamma_i \mid \alpha_0^{(t)}, \alpha_1^{(t)}, d_i, \text{ data }) = \Pr(\gamma_i = 1 \mid d_i, \alpha_0^{(t)}, \alpha_1^{(t)}, \text{ data }) = \frac{\pi_i^{(t)} \mathrm{BF}_i}{(1 - \pi_i^{(t)}) + \pi_i^{(t)} \mathrm{BF}_i} \tag{S11}$$

In the $t$-th M step, we maximize the complete data log likelihood with respect to $(\alpha_0, \alpha_1)$. The maximization procedure is equivalent to fitting a logistic regression or computing from a $2 \times 2$ contingency table with fractional cell counts. Specifically, let

$$\begin{aligned} C_{00}^{(t)} &= \Sigma_{i=1}^m \Big(1 - \mathrm{E}(\gamma_i \mid \alpha_0^{(t)}, \alpha_1^{(t)}, d_i, \text{ data })\Big) \mathbf{1}(d_i = 0) \\ C_{10}^{(t)} &= \Sigma_{i=1}^m \mathrm{E}(\gamma_i \mid \alpha_0^{(t)}, \alpha_1^{(t)}, d_i, \text{ data }) \mathbf{1}(d_i = 0) \\ C_{01}^{(t)} &= \Sigma_{i=1}^m \Big(1 - \mathrm{E}(\gamma_i | \alpha_0^{(t)}, \alpha_1^{(t)}, d_i, \text{ data })\Big) \mathbf{1}(d_i = 1) \\ C_{11}^{(t)} &= \Sigma_{i=1}^m \mathrm{E}(\gamma_i \mid \alpha_0^{(t)}, \alpha_1^{(t)}, d_i, \text{ data }) \mathbf{1}(d_i = 1). \end{aligned} \tag{S12}$$

It follows that

$$\begin{aligned} \hat{\alpha}_0^{(t+1)} &= \log\left[\frac{C_{10}^{(t)}}{C_{00}^{(t)}}\right] \\ \hat{\alpha}_1^{(t+1)} &= \log\left[\frac{C_{00}^{(t)} C_{11}^{(t)}}{C_{10}^{(t)} C_{01}^{(t)}}\right], \end{aligned} \tag{S13}$$

We iterate between E and M steps until the algorithm converges.
